# Computational and Image Processing Methods for Analysis and Automation of Anatomical Alignment and Joint Spacing in Reconstructive Surgery

**DOI:** 10.1101/2021.04.04.438341

**Authors:** Usamah N. Chaudhary, Cambre N. Kelly, Benjamin R. Wesorick, Cameron M. Reese, Ken Gall, Samuel B. Adams, Guillermo Sapiro, J. Matias Di Martino

## Abstract

**Purpose:** Reconstructive surgeries to treat a number of musculoskeletal conditions, from arthritis to severe trauma, involve implant placement and reconstructive planning components. Anatomically matched 3D printed implants are becoming increasingly patient-specific; however the preoperative planning and design process requires several hours of manual effort from highly trained engineers and clinicians. Our work mitigates this problem by proposing algorithms for the automatic re-alignment of unhealthy anatomies, leading to more efficient, affordable, and scalable treatment solutions.

**Methods:** Our solution combines global alignment techniques such as iterative closest points (ICP) with novel joint space refinement algorithms. The latter is achieved by a low dimensional characterization of the joint space, computed from the distribution of the distance between adjacent points in a joint.

**Results:** Experimental validation is presented on real clinical data from human subjects. Compared with ground truth healthy anatomies, our algorithms can reduce misalignment errors by 22% in translation and 19% in rotation for the full foot-and-ankle and 37% in translation and 39% in rotation for the hind-foot only, achieving a performance comparable to expert technicians.

**Conclusion:** Our methods and histogram-based metric allow for automatic and unsupervised alignment of anatomies, a major step toward a fully automated and data driven re-positioning, designing, and diagnosing tool.

## 1 Introduction

Orthopaedic reconstructive surgery seeks to preserve anatomical functionality in cases of musculoskeletal defects. Surgical intervention to treat these conditions often involves repair and realignment by placement of an implant. Orthopaedic surgeries such as hip or knee replacements, as well as replacement of large missing segments or even entire bones with 3D printed implants, are becoming increasingly patient-specific in their planning, design, and placement [1–5]. For all of these reconstructive surgeries, healthy postoperative joint alignment and spacing are critical to restoring normal function, and this is usually achieved by placing articulating components and fixation devices such as screws, rods, or spacers [6–8].

With increasing emphasis on preoperative and patient specific surgical plans for such reconstructions, computed tomography (CT) scans are often required to visualize the anatomy. Since most reconstructive cases involve patients unable to place weight on the affected area, the CT scans obtained are often in a supine (non-weight bearing) position. This presents a major challenge in surgical planning as surgeons and design engineers must design anatomically matched implants with consideration of the weight bearing position, the importance of which has been noted in recent studies [9]. To circumvent this, design engineers manually realign the patient’s anatomy to a theorized weight bearing position with affected anatomy removed. The 3D printed implant must be appropriately designed, a very time-intensive process with the potential for significant inter-engineer variability. This manual process involves the engineer considering all articulations in the anatomy of interest to ensure healthy joint spacing when realigning the bones.

Preoperative planning for reconstructive surgery includes consideration of both joint alignments and articulations. Articulating joints between bones in the human body are lined with hyaline cartilage, an elastic connective tissue that covers the joint surfaces and makes up a majority of the joint space [10]. Articular cartilage enables a range of motion at the joint, as well as transmission of loads [11]; however, it is not visualized on CT, adding to the difficulty of the manual task. Damage or pathological changes to articular cartilage can result in reduction or enlargement of joint spacing, and is the cause of, or is comorbid with several pathologies including various forms of arthritis, trauma, neoplastic growth of bone or cartilaginous tissue, and others [12].

In the present work, we propose robust algorithms to analyze joint spacing and automate the alignment of pathological anatomies. We propose a joint sanity measure to optimize alignment based on the distribution of the point distances between adjacent anatomies. We show this metric can be effectively optimized, leading to different numerical alignment algorithms. We validate the proposed methods with real data from foot-and-ankle anatomies and compare our results with those obtained manually by trained experts. Our main contributions can be summarized as follows: (i) We design a histogram-based metric that can effectively characterize the joint’s sanity while being nearly invariant to the many healthy articulations of a given joint; (ii) Based on our histogram-based metric, we propose a practical solution for the automatic re-alignment of anatomy; (iii) The proposed framework performs comparable to manual alignment by experts, resulting in the first ever scalable solution for reconstructive surgery of musculoskeletal conditions.

### 1.1 Related Work

The use of computerized methods to create implants and analyze joint spacing is a developing field, with most current methods focusing on the design of custom implants [13–15]. Harrysson et al. were able to create significantly more patient-specific knee replacement parts than conventional generalized molds, leading to improved stress distribution and fewer postoperative gait changes. Healthy joint spacing was factored into the implants’ design; however, as noted by the authors, the manual design process is very time-intensive and remains an open problem. This presents a common problem in reconstructive surgery: the pre-surgical planning stage requires a significant amount of manual intervention.

Current methods on analyzing joint spacing are often not highly generalizable and focus on a single characteristic of the joint, such as average spacing, rather than accounting for all the joint complexities and kinematics. Imai et al. were able to calculate joint spacing in the tibiotalar joint and compared spacing width at varying degrees of plantarflexion and dorsiflexion. However, they present a single distance value averaged across the joint, which, although useful in comparing overall spacing, lacks proper analysis of the joint spacing, nor can it be optimized in an automated alignment process. [16]. Siegler et al. presented distance maps of surface-to-surface spacing which showed significant differences in tibiotalar and subtalar joint spacing from neutral to extreme positions in varying triplanar rotations. However, this study was not done under axial loading conditions and did not reproduce kinematics during functional activities such as walking. [17]

Lalone et al. analyzed the joint spacing of the ulna-humerus joint at varying degrees of elbow extension and flexion to map the changes in the bones’ proximity. Their work is, to the best of our knowledge, the closest to ours. Lalone et al. were able to map the distance between bones successfully; however, manual, pre-defined distances were used to consider spacing as high proximity (< 0.5*mm*), medium proximity (< 1.5*mm*), low proximity (< 2.5*mm*) and ultra-low proximity (< 3.5*mm*). Although this method is sound for generating proximity maps for visualization, comparison to the spacing information within a known healthy joint allows for better analysis [18, 19]. A difference between our work and previous work is that we propose a histogram-based joint measure, which can be minimized (using pre-established healthy templates) to achieve automatic anatomical realignment.

Moreover, we show that the proposed metric better captures the condition of a joint when compared to absolute translation and rotation misalignment. This is particularly true in joints that allow the bones to rotate with respect to each other along the joint axis, such as the tibiotalar joint. In these conditions, aligned anatomies are represented by a manifold of solutions; which we explicitly address in our methods, in contrast to previous literature.

## 2 Methods

All parts of the human skeleton, such as the foot-and-ankle, hand, or knee, are articulated and positioned in a particular manner enabling them to achieve their anatomical functions. Moreover, there is a specific arrangement between joints, particularly in the extremities where one bone can have multiple articulations with more than one adjacent bone. In this section, we start by describing and defining alignment into a suitable mathematical framework, and then we focus on the problem of automatic anatomical realignment.

### 2.1 Definitions and notation

In the present work, we focus on foot anatomies that have been generated by 2D image segmentation of CT scans followed by 3D reconstruction of each component (bone). We consider the steps of 3D reconstruction and segmentation pre-processing that are out of the scope of this work (see for example [20–24] for techniques to perform such steps), hence we consider the 28 bones of the foot-and-ankle as the input set ℐ = {𝒫_1_, …, 𝒫_28_} with 𝒫_*i*_ = {***x***_*j*_ *∈ ℝ*^3^, *j* = 1, …, *n*_*i*_}. *n*_*i*_ represents the number of 3D points in the cloud associated with the bone *i*. Some bones could be partially or completely missing for unhealthy anatomies, in which case the set 𝒫 associated with it is defined as the empty set. Notice that in the definition above, we used a bold font to represent a 3D vector; we will keep this convention from now on, ***x*** = (*x*_1_, *x*_2_, *x*_3_) and *x*_*k*_ represent a vector and the value of its *k*^*th*^ coordinate respectively.

For a point ***x*** in the element 𝒫_*i*_, we define its distance to the element 𝒫_*j*_ as, 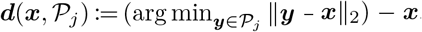, meaning, ***d*** represents the vector that connects ***x*** to its closest neighbor in 𝒫_*j*_. As discussed earlier, the relative position between bones and the configuration of the joints is one of the most critical aspects to achieve a healthy alignment and we introduce two definitions to address this: 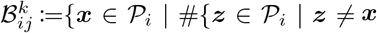 and 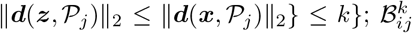 represents the set of *k* closest points to 𝒫_*j*_ vectors in 𝒫_*i*_. Finally, the joint space between 𝒫_*i*_ and 𝒫_*j*_, is characterized by the set 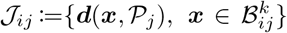. Figure 1 illustrates the definitions introduced above, as illustrated in Figure 1(b) the problem can be represented as a graph of components 𝒫_*i*_, with interconnections characterized by joints represented by 𝒥_*ij*_.

**Fig. 1.**
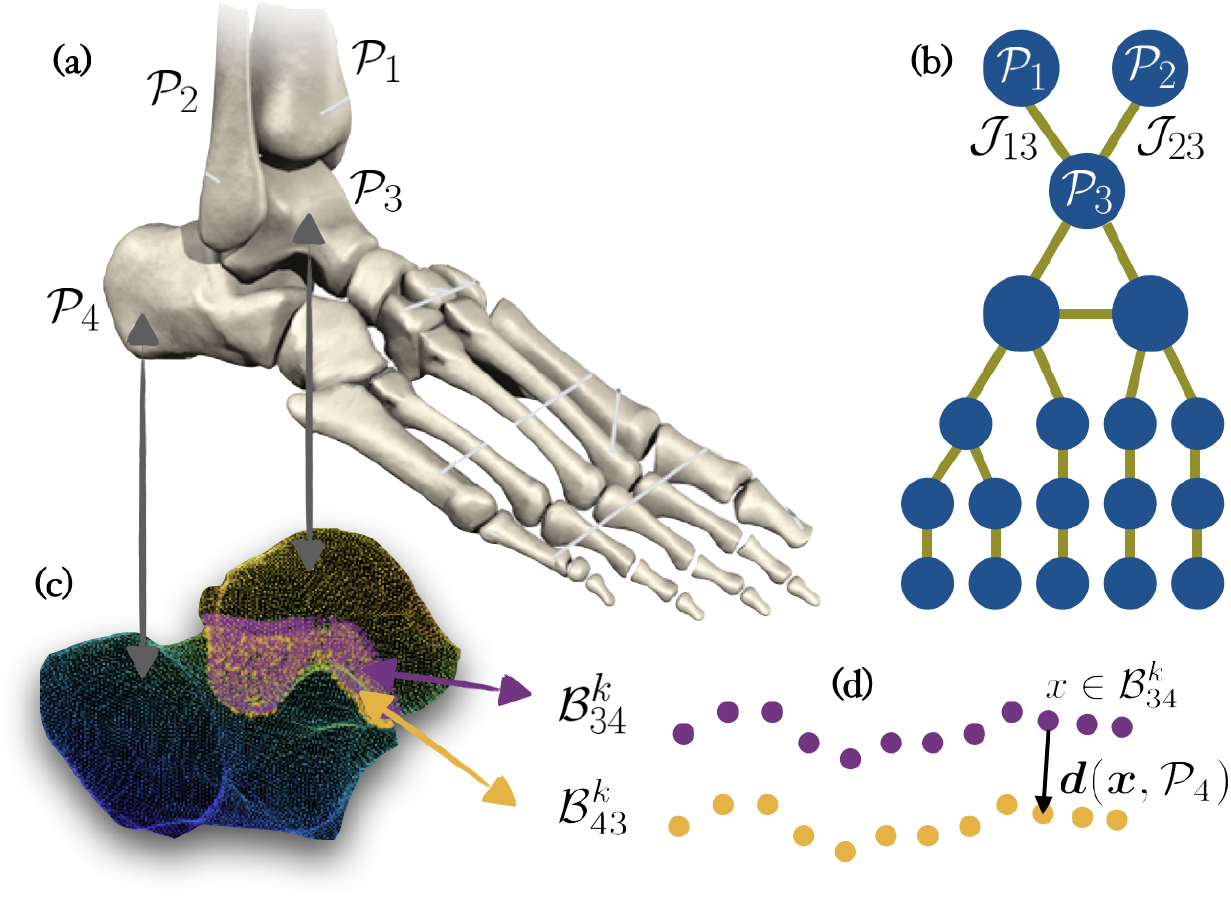
Problem definition and notation. (a) A healthy input anatomy, each part (bone) is represented by the set of 3D points 𝒫_*i*_. (b) An abstract (graph) representation of the components and their joints interconnection. (c) Space between the talus and the calcaneus, and the set of *k* closest points represented by 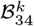 and 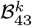, respectively. (d) Distance between a point *x* ε 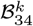 and its adjacent bone represented by 𝒫_4_.

### 2.2 Analysis of the joint space

Given the sets of 3D points 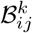, we can computationally analyze the joint space and design metrics for determining the health of a joint. Quantitative determination of health is the first step in the automated anatomical realignment. It is also a useful metric on its own that can be used in the diagnosis of joint spacing diseases such as osteoarthritis (OA) as previously discussed. The health of the joint between 𝒫_*i*_ and 𝒫_*j*_ lies in the distribution of the distances of the one to one correspondences used to generate the two sets 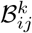 and 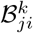, given by the set 𝒥_*ij*_. This distribution for a healthy joint contains all the information about distances, uniformity, and density needed to determine a health metric. We define the histogram *h*_*ij*_ :=[#{‖ ***d***_*q*_ ‖ < *r*_1_}, #{*r*_1_ ≤ ‖ ***d***_*q*_ ‖ *< r*_2_}, …, #{*r*_*m*_ ≤ ‖ ***d***_*q*_ ‖}], with ***d***_*q*_ in 𝒥_*ij*_, and *r*_1_, …, *r*_*m*_ representing the range of the histogram (bins); these values are manually tuned based on the visual inspection of several healthy anatomies.

We hypothesize that the histogram of the norm of the distances between neighbors in a joint can be used as a low dimensional representation to describe the joint space’s morphology. Hence, it can be used as an objective function for optimizing automatic re-alignment, as we show next. As illustrated in Figure 3 (f), a healthy joint distribution contains a large density of points in a very narrow window indicating a uniform gap space at a specific distance. Any abnormalities in the joint such as joint space narrowing will be seen visually on this distribution as shown in the overlaid histogram in figures 2 and 3.

**Fig. 2.**
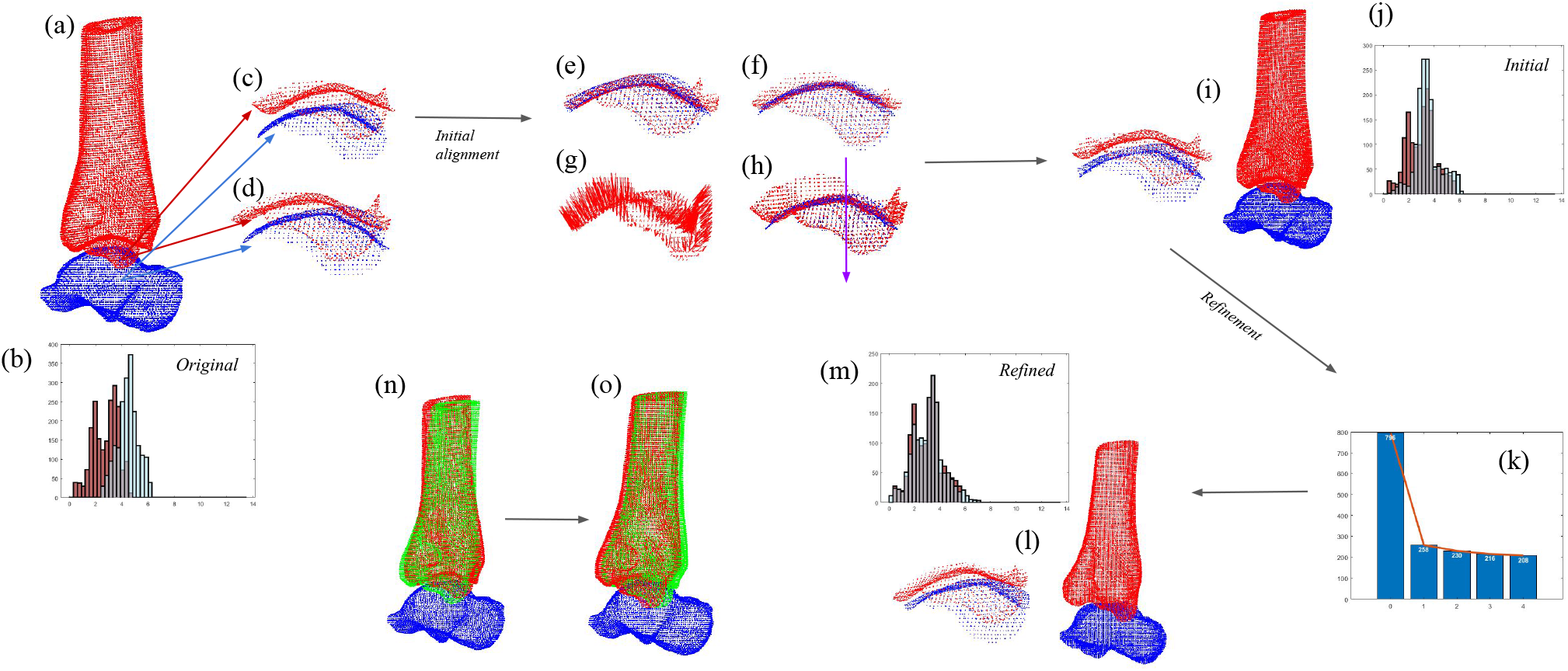
Automated alignment method. (a) Pathological arrangement of the tibia (red) and talus (blue). (b) Histogram of the absolute distance between corresponding pairs in the adjacent surfaces (*h*_*ij*_ and 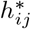 defined above), the distribution of the input (unhealthy) anatomy is illustrated i in light blue, and the distribution of the ground truth (healthy) template in maroon. (c) Articulating surface of the pathological (input) example. (d) Ground truth (healthy) target alignment. (e-h) Evolution of the initial alignment: (e) and (f) illustrate two steps of ICP mating, (g) shows the vector directions between the two surfaces, and (h) the average vector direction. Similar to (a-c), (i-j) shows the aligned anatomy and its corresponding histogram after the initial alignment. During the refinement step, the distance between the target histogram and the empirical histogram is minimized using a multi-grid optimization, the evolution of the residual is illustrated in (k). (m-l) Final anatomy aligned (and its associated histogram of distances in light blue, as before, the template histogram is illustrated in maroon). (n) and (o) compares the starting position (n) and the final result (o), in green we overlay the ground truth healthy anatomy.

**Fig. 3.**
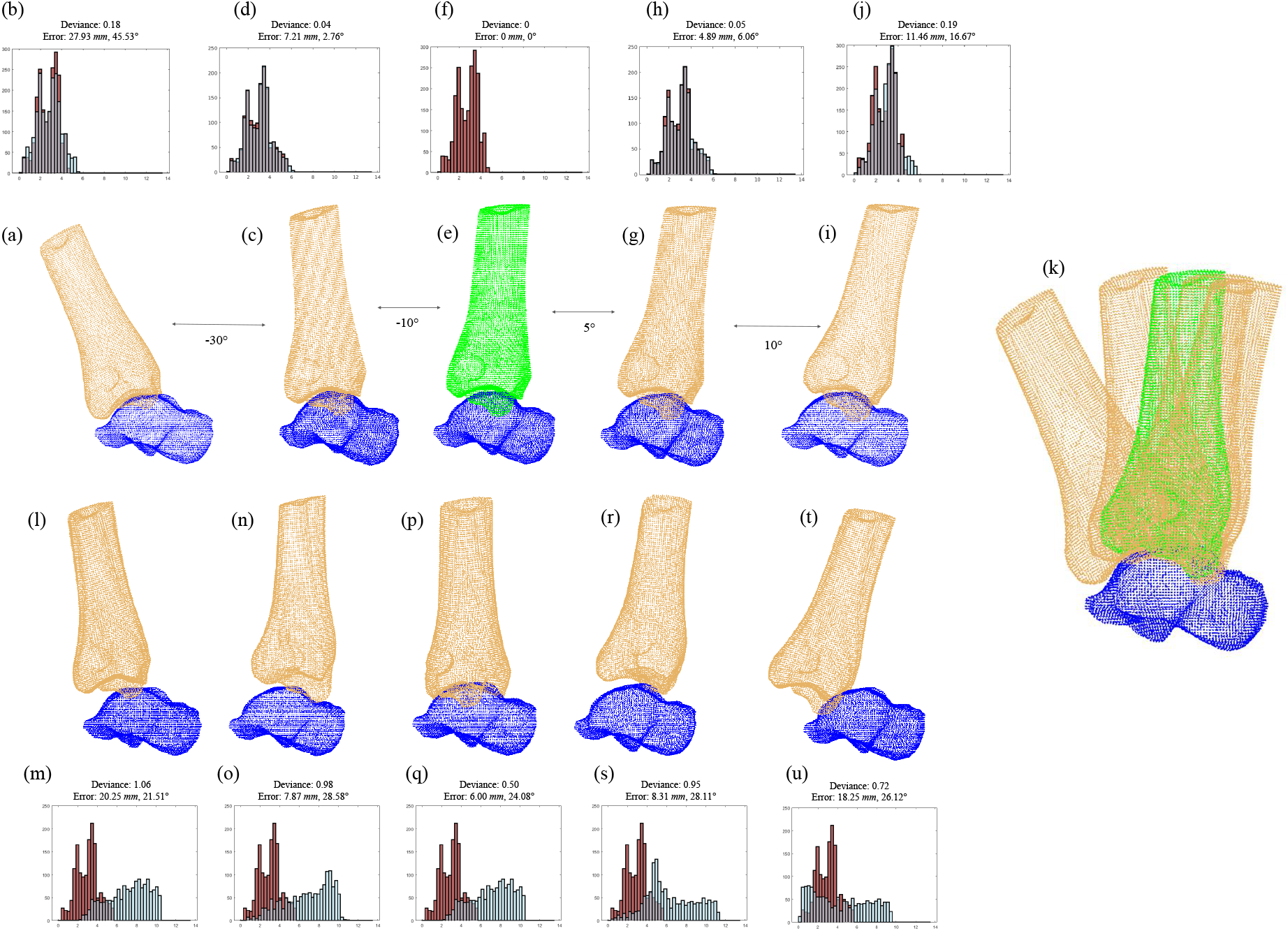
Deviance value robustness. (a) 30 degrees of plantarflexion of tibia-talus joint along with its distribution in blue overlayed with the healthy distribution in maroon shown in (b). (c) 10 degrees of plantarflexion along with distribution in (d). (e) original position of the scan along with distribution in (f). (g) 5 degrees of dorsiflexion and distribution in (h). (i) 10 degrees of dorsiflexion and distribution in (j). (k) illustrates (a), (c), (e), (g), and (i) all overlayed. (l) through (u) shows five random translations and rotations in the same space as (a) through (j).

### 2.3 Automatic anatomical re-alignment

Automated anatomical alignment is performed in two main steps. First, we extract the articulating surfaces of both bones as described in Subsection 2.1; since these surfaces are complementary, we perform a series of heuristic steps to align and pull the anatomies apart to a favorable position. Second, we perform a small local search to identify the optimal positioning. This local search allows the algorithm to adjust for small changes in how the bones align outside of just the two anatomies fitting together, such as from the pulls of various ligaments and tendons on the bones.

#### Initial alignment

We first extract an initial estimation of the articulating surfaces between anatomies 𝒫_*i*_ and 𝒫_*j*_, giving 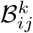 and 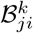 respectively; then we perform an adaptation of the Iterative Closest Points (ICP) registration algorithm [25] between the set of points. This registration creates a mate between the articulating (complementary) surfaces of the two anatomies. After this initial mating, we recalculate the final boundary sets,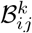 and 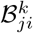, and repeat the ICP (shown in Figure 2 (e) and (f)). We empirically observed that this process leads to a robust calculation of the articulation surfaces, even under large initial anatomy deformations (further discussion is presented in Section 3).

Finally, to achieve anatomically sound joint spacing, we pull the bones apart from the mated position by increasing the distance between the bones. To that end, we move one of the bones along the direction given by 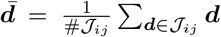, shown in purple in Figure 2 (h). At each iteration, we optimize the positioning by minimizing the histogram distance between the sample *h*_*ij*_ and a healthy distribution 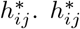 represents the template histogram distribution for a given joint. We can obtain such a template from (a) data from previous (healthy) patients, (b) manually aligned anatomies, or (c) the contralateral (opposite side of body) scan of the same patient if available. The *L*2 distance between the empirical distribution *h*_*ij*_ and the template distribution 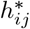, also referred to as the deviance henceforth, is used to assess the registration error. Typically a distance up to 5*mm* is sufficient to check as all joint gaps in the human body are within that range [26–29]. As exemplified in Figure 2 (e) through (h), this method mates the complementary articulating surfaces together and correctly identifies the direction of movement to pull the anatomies apart.

#### Positioning refinement

After the initial re-positioning, we optimize the relative position of the bones, by continuing to minimize the *L*2 distance between *h*_*ij*_ and 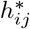. A bone is a rigid body, hence it’s position can be fully represented by six parameters, three to establish the location of its center (*x, y, z*), and three its orientation (*θ*_1_, *θ*_2_, *θ*_3_). We continue optimizing the alignment by testing in the six-dimensional parameter space, the set of values (*x, y, z, θ*_1_, *θ*_2_, *θ*_3_) that minimize the *L*2 distance between *h*_*ij*_ and 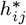.

Since an exhaustive search in a fine six-dimensional grid can be computationally expensive, we optimize these parameters in a multi-scale fashion. In each iteration step, we test all the possible translations and rotations on a given resolution. Once the optimal value is obtained, we refine the parameters resolution around this neighborhood and optimize within this new (refined) space. This process is repeated until the distance between the histograms is below a pre-defined threshold. Figure 2 illustrates the evolution of the deviance between the histograms as the multi-grid optimization iterated Figure 2-(k), the target (maroon) and final (light blue) histograms are illustrated in Figure 2-(m).

## 3 Results

### Data

CT scans of both healthy and unhealthy foot-and-ankle anatomies were retrospectively obtained from previous surgical cases. The CT scans were manually segmented (although multiple techniques exist to make this automatic, being this out of the scope of this paper), and the CT scan slices were used to generate 3D scans represented by a set of point clouds. Our primary data set includes 9 healthy weight bearing scans and 9 pathological scans. The foot-and-ankle region contains 28 bones and 33 unique joints. The 3D representation of the anatomy has an average resolution of 98, 955 3D points distributed as 22, 466 for the tibia, 9, 880 for the talus, 4, 236 for the medial cuneiform, 2, 042 for the intermediate cuneiform, 2, 618 for the lateral cuneiform, 4, 772 for the navicular, 15, 881 for the calcaneus, 6, 511 for the first metatarsal, 4, 580 for the second metatarsal, 4, 074 for the third metatarsal, 3, 946 for the fourth metatarsal, and 4.379 for the fifth metatarsal. Our methods and results exclude discussion and calculation on the phalanges and fibula as they are less relevant to most studied surgical cases.

### Tuning model parameters

After a visual inspection of the healthy samples, we set the number of points (*k*) in the 3D model that defines the joint sets 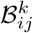. The empirical values obtained are as follows: 12.4% and 18.2% for the Tibia-Talus joint, 13.0% and 18.3% for the Calcaneus-Talus joint, 10.1% and 21.8% for the Navicular-Talus joint, 5.2% and 17.7% for the Calcaneus-Cuboid joint, 15.5% and 13.8% for the Medical Cuneiform-Navicular joint, 17.2% and 7.9% for the Intermediate Cuneiform-Navicular joint, 11.1% and 5.9% for the Lateral Cuneiform-Navicular joint, 13.0% and 22.2% for the 1st Metatarsal-Medical Cuneiform joint, 10.5% and 21.5% for the 2nd Metatarsal-Intermediate Cuneiform joint, 12.6% and 18.5% for the 3rd Metatarsal-Lateral Cuneiform joint, 4.7% and 4.4% for the 4th Metatarsal-Cuboid joint, 4.3% and 4.4% for the 5th Metatarsal-Cuboid joint. All the values are provided as a percentage of the number of points in the articulation in the first and second bone, respectively, for each joint. (We provide relative values since the absolute number of points in a joint will be highly dependant on the scanning resolution.) The joints histograms *h*_*ij*_ are defined on the range 0 − 13.5*mm*. Multi-scale joint refinement search is performed in four nested resolutions defined by [±0.5*mm*, ±5°], [±0.1*mm*, ±1°], [±0.02*mm*, ±0.2°], and [±0.004*mm*, ±0.04°]. Locally, each scale is centered around the optimum value in the previous coarser search.

### Automatic joint re-alignment

To quantitatively test the proposed re-alignment algorithms, we generated misaligned examples starting from the samples of healthy anatomies. Table 1 shows, for each joint, the initial error, the error after alignment, and the relative improvement. Across the 9 healthy cases, random deformations were applied to the 12 major joints accounting for 108 2-body test cases (individual results of each test case are shown in Tables S1-S9). In Table 1 we report the norm of the translation and rotation vector along with a normalized *L*2 deviance value between the cases and the ground truth distribution. Across all cases, the average applied deformation was (2.03 *mm*, 14.32°) from which the algorithm was able to achieve an error of (1.55 *mm*, 11.01°), representing an improvement in absolute error of 22.14% in translation and 19.05% in rotation. The normalized deviance value was attained by converting the distribution of points into a probability distribution such that ∫ *h*_*ij*_ = 1. The *L*2 distance between probability distributions is reported in Table 1. These values range from 0, representing a perfect overlap of the distributions, to 2, representing a lack of any overlap, the normalized deviance value across all cases and joints improved by 86.0%. If excluding the metatarsals, which can have a high degree of displacement from healthy plantarflexion or dorsiflexion and are often not instrumental to surgery as the majority of reconstructive cases involve the hind foot region, the improvement is 37.22% in translation and 39.42% in rotation, with an 86.15% drop in deviance values.

**Table 1.** Autoamted alignment results. Using a pair of healthy and unhealthy samples, we report the original and final translation ‖ (*x, y, z*) ‖ and rotation ‖ (*θ*_1_, *θ*_2_, *θ*_3_) ‖ error, along with the normalized *L*2 histogram deviances. Each row represents the average results across the 9 patient cases, with the average for all joint in each case shown in the final row.

| Joint | Average Applied Deformation | Average Algorithmic Error | Relative Improvement |
| --- | --- | --- | --- |
| Anatomy 1 - Anatomy 2 | Translation (mm), Rotation (°), histogram $L2$ Deviance | | |
| Tibia - Talus | 1.91 mm, 12.62 °, 0.56 | 1.22 mm, 7.27 °, 0.06 | 36.23%, 42.46%, 88.88% |
| Calcaneus - Talus | 2.00 mm, 13.12 °, 0.87 | 1.42 mm, 6.67 °, 0.11 | 29.01%, 49.15%, 88.14% |
| Navicular - Talus | 1.92 mm, 26.80 °, 0.83 | 0.97 mm, 9.10 °, 0.07 | 49.67%, 66.03%, 90.92% |
| Calcaneus - Cuboid | 2.39 mm, 13.93 °, 0.61 | 1.06 mm, 10.73 °, 0.16 | 55.65%, 22.97%, 73.67% |
| Med. Cuneiform - Navicular | 1.83 mm, 14.45 °, 0.75 | 1.62 mm, 8.42 °, 0.12 | 11.49%, 41.74%, 81.39% |
| Int. Cuneiform - Navicular | 2.30 mm, 12.50 °, 1.28 | 1.28 mm, 8.97 °, 0.04 | 44.26%, 28.23%, 85.32% |
| Lat. Cuneiform - Navicular | 1.99 mm, 14.27 °, 1.25 | 1.31 mm, 10.65 °, 0.05 | 34.23%, 25.36%, 94.65% |
| 1st Metatarsal - Med. Cuneiform | 1.86 mm, 13.06 °, 0.72 | 1.23 mm, 11.14 °, 0.10 | 33.93%, 14.66%, 85.35% |
| 2nd Metatarsal - Int. Cuneiform | 2.51 mm, 11.99 °, 0.84 | 1.46 mm, 13.81 °, 0.09 | 41.93%, -15.23%, 86.97% |
| 3rd Metatarsal - Lat. Cuneiform | 1.64 mm, 13.02 °, 0.74 | 1.79 mm, 12.90 °, 0.11 | -9.19%, 0.93%, 83.51% |
| 4th Metatarsal - Cuboid | 1.92 mm, 13.60 °, 1.20 | 2.06 mm, 18.37 °, 0.20 | -7.04%, -35.10%, 81.58% |
| 5th Metatarsal - Cuboid | 2.05 mm, 12.47 °, 1.08 | 3.16 mm, 14.04 °, 0.06 | -54.53%, -12.63%, 91.51% |
| **Average - Full Foot-and-Ankle** | **2.03 mm, 14.32 °, 0.89** | **1.55 mm, 11.01 °, 0.10** | **22.14%, 19.05%, 86.0%** |
| **Average - Hind-Foot** | **2.05 mm, 15.38 °, 0.88** | **1.27 mm, 8.83 °, 0.09** | **37.22%, 39.42%, 86.15%** |

### A healthy anatomy can be represented by multiple solutions

One of the main challenges when reporting alignment errors is that a healthy anatomy is represented by a continuum of solutions rather than a fixed body. The foot has many healthy articulations under weight bearing, partial weight bearing, and non-weight bearing movements compatible with normal range of motion, all of which represent healthy anatomies. We illustrate this in Figure 3(a)-(j), where a normal movement of the foot is represented with its associated relative positions for the tibia and talus. Hence, measuring the relative position between anatomies is not a robust metric of anatomical conditions. On the other hand, the proposed histogram-based metric is a practical metric to assess the state of the anatomy and joints. This is illustrated in Figure 3. On top, (a)-(j), we show a healthy movement, and as we can see, the histogram of distances associated with each sample remains almost invariant. In contrast, we present (l)-(u) cases with a deformation of similar amplitude but with respect to a random axis; the later models an unhealthy misalignment. As we can see, the proposed histogram-based metric is more sensitive to the examples’ healthiness and provides a more meaningful measure of error than the deviance of absolute angles.

### Comparison to expert manual alignment

In order to frame the strength of our approach in the context of real world clinical challenges and standard of care manual alignment, we compared our algorithmic process to 5 trained expert design engineers at restor3d. We applied a series of joint spacing deformations to a healthy scan and compared manual aligned joint deviance values to algorithmic deviance values, shown in Table 2. The average applied deformation was a normalized deviance of 1.57. The five engineers (agnostic to the ground truth and the applied deformations) achieved an average deviance of 0.85, representing a 42.90% improvement, in approximately 1 hour each. In Table 2 “Algo with Contralateral,” we report the results from using the contralateral scan for the patient for the global ICP, while in the “Algo with Non-Contralateral” column, we report the results from a non-contralateral scan to simulate results from situations in which the contralateral scan may not be available. Both achieve similar results at 94.23% and 91.32% improvements, respectively.

**Table 2.** Comparison to expert engineer manual alignments. The deformed column values are the normalized deviance values for applied deformation for each joint. The Engineer Average column values are the average normalized deviance values achieve by the 5 engineers along with percent improvements in parenthesis. The Algo with Contralateral and Algo with Non-Contralteral columns display values achieved by our methods with an initialization with a contralateral and non-contralateral scan, respectively. The Navicular-Talus joint is not included as it was given in a non-deformed state to the engineers.

| Joint | Applied Deformation | Engineer Average | Algo with Contralateral | Algo with Non-Contralateral |
| --- | --- | --- | --- | --- |
| Tibia - Talus | 1.87 | 0.69 (63.25%) | 0.07 (96.26%) | 0.09 (95.34%) |
| Calcaneus - Talus | 1.55 | 0.67 (56.61%) | 0.09 (94.05%) | 0.06 (95.86%) |
| Calcaneus - Cuboid | 1.86 | 0.58 (68.86%) | 0.05 (97.15%) | 0.18 (90.35%) |
| Med. Cuneiform - Navicular | 0.89 | 0.54 (39.47%) | 0.13 (84.98%) | 0.14 (84.18%) |
| Int. Cuneiform - Navicular | 1.96 | 1.31 (33.11%) | 0.02 (98.91%) | 0.11 (94.56%) |
| Lat. Cuneiform - Navicular | 1.27 | 1.32 (-3.90%) | 0.08 (94.02%) | 0.16 (87.61%) |
| **Average** | **1.57** | **0.85 (42.90%)** | **0.07 (94.23%)** | **0.12 (91.32%)** |

## 4 Discussion

We introduce a metric for analyzing joint spacing between two bodies and automatic alignment methods. The proposed techniques achieve an 86.0% decrease in alignment deviance and an improvement of 22.14%/19.05% in translation/rotation positioning error. When considering only the hind foot, our methods perform even better at 37.22%/39.42% for improvement in the translation/rotation alignment, along with a 86.15% decrease in histogram deviance. Compared to expert engineers (0.85 deviance), our methods (0.12 deviance) can attain improved joint spacing.

Our proposed methods are a first step toward helping automate an expensive and time-consuming manual process. Much of the current literature on 3D printed implants and reconstructive surgery lacks robust methods for joint space analysis. Furthermore, relative position is a non-robust metric for anatomy alignment since articulations such as the tibia-talus enable the (healthy) rotation with respect to a specific axis. Thus a CT scan, weight-bearing or not, is merely a single solution of one of these possible arrangements. Therefore, the errors shown in Table 1 should be taken contingently as any alignment, even with healthy joint spacing and weight-bearing position, that is not the exact atlas position will be penalized. We overcome this issue and propose a novel and useful metric that can be used to assess and correct anatomical misalignment. To the best of our knowledge, this is a new metric, with further clinical implications beyond alignments.

Although our results already present marked improvement based on absolute error alone (both for the translation and rotation components), they are more impressive in the context of the histogram deviance improvement. Across the 108 tested cases, our algorithm was able to reduce deviance values by 86.00%. All metatarsals show improvement in deviance values; however, the 2nd, 3rd, 4th, and 5th metatarsals show worsened absolute error; this is likely due to a large healthy range of motion for these bones, as previously discussed.

Our empirical results strengthen our hypothesis that a histogram-based metric can effectively characterize the joint’s sanity while being nearly invariant to various healthy articulations of a given joint. In addition, the proposed automatic realignment method effectively minimized the proposed deviance metric, leading to better joint spacing and healthier anatomical solutions.

### Limitations

In the comparison results presented in Table 2, the 5 expert engineers were given the task of performing a global alignment with healthy joint spacing while our algorithms were driving toward only healthy joint spacing. Engineers may have sacrificed joint spacing health on particular joints to achieve better global alignments. In that sense, these results validate that our method achieves a joint alignment accepted by human experts and below the error margins they consider defines a successful joint alignment. However, these results do not indicate our method outperforms humans since they were solving a harder task. Thus, global positioning must be ultimately considered during alignment, as alignment of one joint to the detriment of another is not valuable. In the present work, the methods and experiments explore the two-body problem and focus on individual joints. In that sense, our work does not solve the global alignment of all the anatomical components simultaneously. Rather, we focus on the joint alignment given two adjacent bones. Solving global anatomical alignment is important and will be addressed in future work. We believe the methods proposed here will be critical building blocks of the final global alignment solution.

### Future work

Future work includes, in addition to the study of the multi-body problem discussed above, the collection of a larger database of anatomies and the study of theoretical guarantees for the convergence of the proposed methods. Collecting an extensive database of healthy and unhealthy anatomies is crucial to obtain a representative and diverse sample to ensure that templates are available to fit any patient of any age, sex, and demographic. Moreover, an extensive database would enable us to explore entirely data driven approaches to leverage the recent advances in deep learning.

## 5 Conclusions

We presented automatic algorithms for joint realignment and the automatic registration of unhealthy anatomies. Posing the problem into a proper mathematical framework allows us to set the algorithmic foundations for a broader objective: a fully automated and data driven re-positioning, designing, and diagnosing tool. We validated the proposed ideas with real clinical data from healthy and unhealthy anatomy from patient CT scans. Experiments suggest that the proposed histogram-based metric is more suitable for assessing anatomical conditions than absolute translation and rotation error. More importantly, the proposed metric can be efficiently optimized via multi-grid search algorithms, leading to a fully automatic and unsupervised joint alignment tool.

## Acknowledgments

The work of GS and JMDM is partially supported by the National Science Foundation, the Department of Defense, and gifts from Amazon Web Services, Microsoft, Google, and Cisco.

