## Supplementary Tables S1-9 for "Computational and Image Processing Methods for Analysis and Automation of Anatomical Alignment and Joint Spacing in Reconstructive Surgery"

**Noname manuscript No.**
(will be inserted by the editor)

### SUPPLEMENTARY MATERIAL: Computational and Image Processing Methods for Analysis and Automation of Anatomical Alignment and Joint Spacing in Reconstructive Surgery

**Usamah N. Chaudhary* · Cambre N. Kelly · Benjamin R. Wesorick · Cameron M. Reese · Ken Gall · Samuel B. Adams · Guillermo Sapiro · J. Matias Di Martino**

Received: date / Accepted: date

#### Supplementary Material

U.N. Chaudhary
UT Southwestern Medical Center
Dallas, TX 76126, USA

*Corresponding author

C.N. Kelly, B.R. Wesorick, C.M. Reese
restor3d, Inc.
311 West Corporation St. Durham, NC 27701, USA

K. Gall
Duke University, Pratt School of Engineering
Durham, NC, 27701, USA

S.B. Adams
Duke University Medical Center
Durham, NC 27705, USA

G. Sapiro, J.M.D. Martino
Duke University
Durham, NC 27705, USA

| Case #1 | | | | | | | | | | | | |
|---|---|---|---|---|---|---|---|---|---|---|---|---|
| | Medial Cuneiform - Navicular | | | | | | 4th Metatarsal - Cuboid | | | | | |
| Applied Deformation | -1.00 | -2.00 | 0.00 | 5.00 | -9.00 | -7.00 | 0.00 | 1.00 | -1.00 | -6.00 | -4.00 | 2.00 |
| Deformation in PCA | -1.00 | -2.00 | 0.00 | 12.65 | 2.51 | 12.42 | 0.00 | 1.00 | -1.00 | 6.27 | 7.46 | 4.33 |
| Algorithmic Error | -2.23 | 0.12 | 0.43 | 7.86 | 5.55 | 7.99 | -1.89 | -0.16 | 1.86 | 2.91 | 10.16 | 10.20 |
| Deformed Deviance | 1.41 | | | | | | 0.99 | | | | | |
| Algorithmic Deviance | 0.12 | | | | | | 0.05 | | | | | |
| | Intermediate Cuneiform - Navicular | | | | | | 5th Metatarsal - Cuboid | | | | | |
| Applied Deformation | -1.00 | 0.00 | 0.00 | -4.00 | -10.00 | -3.00 | -1.00 | 2.00 | 0.00 | 5.00 | 9.00 | -4.00 |
| Deformation in PCA | -1.00 | 0.00 | 0.00 | 8.50 | 7.16 | 11.05 | -1.00 | 2.00 | 0.00 | 5.06 | 10.50 | 10.07 |
| Algorithmic Error | -0.68 | 0.43 | -0.16 | 4.48 | 1.75 | 4.57 | -1.76 | 0.97 | -0.41 | 10.16 | 6.88 | 11.86 |
| Deformed Deviance | 1.03 | | | | | | 1.68 | | | | | |
| Algorithmic Deviance | 0.01 | | | | | | 0.05 | | | | | |
| | Lateral Cuneiform - Navicular | | | | | | Cuboid - Calcaneus | | | | | |
| Applied Deformation | 1.00 | 2.00 | 1.00 | 1.00 | 9.00 | -9.00 | -2.00 | 2.00 | 0.00 | 5.00 | 0.00 | 1.00 |
| Deformation in PCA | 1.00 | 2.00 | 1.00 | 12.02 | 9.72 | 9.14 | -2.00 | 2.00 | 0.00 | 2.99 | 4.59 | 4.69 |
| Algorithmic Error | 0.15 | 0.75 | -0.59 | 7.48 | 5.90 | 8.86 | -0.09 | 0.17 | -0.20 | 6.24 | 3.41 | 6.04 |
| Deformed Deviance | 1.70 | | | | | | 0.90 | | | | | |
| Algorithmic Deviance | 0.01 | | | | | | 0.11 | | | | | |
| | 1st Metatarsal - Medial Cuneiform | | | | | | Navicular - Talus | | | | | |
| Applied Deformation | 1.00 | 1.00 | 2.00 | 6.00 | -1.00 | 4.00 | 1.00 | 0.00 | 1.00 | -8.00 | -5.00 | 5.00 |
| Deformation in PCA | 1.00 | 1.00 | 2.00 | 7.25 | 3.87 | 6.13 | 1.00 | 0.00 | 1.00 | 4.86 | 9.80 | 10.72 |
| Algorithmic Error | 0.89 | -0.29 | 0.73 | 5.92 | 5.40 | 3.07 | -0.65 | 0.07 | 0.19 | 4.54 | 3.56 | 2.97 |
| Deformed Deviance | 0.97 | | | | | | 0.82 | | | | | |
| Algorithmic Deviance | 0.09 | | | | | | 0.04 | | | | | |
| | 2nd Metatarsal - Intermediate Cuneiform | | | | | | Calcaneus - Talus | | | | | |
| Applied Deformation | -2.00 | 2.00 | -2.00 | 7.00 | -3.00 | -1.00 | 1.00 | -2.00 | -2.00 | 3.00 | 7.00 | -7.00 |
| Deformation in PCA | -2.00 | 2.00 | -2.00 | 7.11 | 7.43 | 3.59 | 1.00 | -2.00 | -2.00 | 4.42 | 9.35 | 10.09 |
| Algorithmic Error | 0.85 | 1.32 | -2.70 | 13.71 | 14.67 | 14.34 | 1.54 | -0.99 | -0.51 | 4.56 | 4.16 | 6.10 |
| Deformed Deviance | 0.82 | | | | | | 0.59 | | | | | |
| Algorithmic Deviance | 0.10 | | | | | | 0.03 | | | | | |
| | 3rd Metatarsal - Lateral Cuneiform | | | | | | Tibia - Talus | | | | | |
| Applied Deformation | 1.00 | 2.00 | 1.00 | -3.00 | -9.00 | -9.00 | 2.00 | 1.00 | -1.00 | 2.00 | 4.00 | 10.00 |
| Deformation in PCA | 1.00 | 2.00 | 1.00 | 5.02 | 11.89 | 12.90 | 2.00 | 1.00 | -1.00 | 4.90 | 10.79 | 10.11 |
| Algorithmic Error | 0.72 | 0.52 | -0.13 | 4.81 | 8.50 | 9.47 | -0.11 | -0.68 | -0.03 | 1.59 | 7.19 | 7.34 |
| Deformed Deviance | 0.79 | | | | | | 0.44 | | | | | |
| Algorithmic Deviance | 0.14 | | | | | | 0.05 | | | | | |

**Table S1** Raw automated alignment results for Case 1. Below each joint, applied random deformation in (x,y,z), applied random deformation in the principle component axes, algorithmic error in principle component axes, normalized deformed deviance, and normalized algorithmic deviance are given. The first three columns for "Applied Deformation" give the applied centroid movement in x, y, and z, respectively, while the final three columns give the applied x, y, and z rotations, respectively. The first three columns in "Deformation in PCA" give the centroid distance between the deformed and ground truth states in x, y, and z, respectively, while the final three columns give the rotations in each principle component axes between the deformed and ground truth states as first major axes, second major axes, and third major axes, respectively. The first three columns in "Algorithmic Error" give the centroid distance between the algorithmic and ground truth states in x, y, and z, respectively, while the final three columns give the rotations in each principle component axes between the algorithmic and ground truth states as first major axes, second major axes, and third major axes, respectively.

| Case #2 | | | | | | | | | | | | |
|---|---|---|---|---|---|---|---|---|---|---|---|---|
| | Medial Cuneiform - Navicular | | | | | | 4th Metatarsal - Cuboid | | | | | |
| Applied Deformation | -2.00 | 0.00 | 0.00 | -9.00 | 5.00 | -3.00 | 0.00 | 0.00 | -2.00 | 9.00 | 4.00 | -1.00 |
| Deformation in PCA | -2.00 | 0.00 | 0.00 | 5.61 | 10.81 | 9.28 | 0.00 | 0.00 | -2.00 | 9.73 | 9.78 | 2.09 |
| Algorithmic Error | -0.46 | 0.59 | -0.75 | 3.67 | 3.52 | 5.05 | -1.73 | 1.04 | -1.80 | 9.99 | 4.08 | 9.65 |
| Deformed Deviance | 0.52 | | | | | | 1.85 | | | | | |
| Algorithmic Deviance | 0.11 | | | | | | 0.14 | | | | | |
| | Intermediate Cuneiform - Navicular | | | | | | 5th Metatarsal - Cuboid | | | | | |
| Applied Deformation | 0.00 | 2.00 | 2.00 | 7.00 | -4.00 | 3.00 | 2.00 | -2.00 | 2.00 | 0.00 | -4.00 | 6.00 |
| Deformation in PCA | 0.00 | 2.00 | 2.00 | 3.62 | 8.41 | 7.82 | 2.00 | -2.00 | 2.00 | 6.64 | 3.90 | 6.68 |
| Algorithmic Error | 0.56 | 0.07 | 0.27 | 0.70 | 1.35 | 1.15 | 0.46 | -1.83 | 2.10 | 6.50 | 15.25 | 16.23 |
| Deformed Deviance | 2.00 | | | | | | 2.00 | | | | | |
| Algorithmic Deviance | 0.03 | | | | | | 0.08 | | | | | |
| | Lateral Cuneiform - Navicular | | | | | | Cuboid - Calcaneus | | | | | |
| Applied Deformation | 0.00 | 0.00 | -2.00 | -6.00 | 8.00 | 0.00 | 2.00 | 2.00 | 2.00 | -3.00 | -6.00 | 8.00 |
| Deformation in PCA | 0.00 | 0.00 | -2.00 | 3.16 | 9.50 | 9.98 | 2.00 | 2.00 | 2.00 | 9.97 | 4.16 | 10.30 |
| Algorithmic Error | -0.65 | -0.32 | -0.10 | 7.50 | 7.18 | 2.67 | 0.65 | -0.42 | 1.36 | 2.52 | 3.01 | 1.72 |
| Deformed Deviance | 1.23 | | | | | | 0.40 | | | | | |
| Algorithmic Deviance | 0.03 | | | | | | 0.06 | | | | | |
| | 1st Metatarsal - Medial Cuneiform | | | | | | Navicular - Talus | | | | | |
| Applied Deformation | 1.00 | 0.00 | 0.00 | 2.00 | 2.00 | -6.00 | 1.00 | 0.00 | -1.00 | 0.00 | -8.00 | 10.00 |
| Deformation in PCA | 1.00 | 0.00 | 0.00 | 6.60 | 0.41 | 6.59 | 1.00 | 0.00 | -1.00 | 12.20 | 3.87 | 12.80 |
| Algorithmic Error | 0.02 | -0.14 | -0.71 | 4.14 | 3.67 | 2.02 | 0.00 | 0.52 | -0.69 | 4.22 | 5.44 | 3.44 |
| Deformed Deviance | 0.65 | | | | | | 0.82 | | | | | |
| Algorithmic Deviance | 0.10 | | | | | | 0.03 | | | | | |
| | 2nd Metatarsal - Intermediate Cuneiform | | | | | | Calcaneus - Talus | | | | | |
| Applied Deformation | 1.00 | -1.00 | -2.00 | -9.00 | 7.00 | 3.00 | 2.00 | -2.00 | -1.00 | 10.00 | -5.00 | 7.00 |
| Deformation in PCA | 1.00 | -1.00 | -2.00 | 8.58 | 10.96 | 8.78 | 2.00 | -2.00 | -1.00 | 8.63 | 12.68 | 9.99 |
| Algorithmic Error | -0.26 | -0.20 | 0.34 | 4.67 | 3.06 | 3.61 | 0.58 | 1.03 | 0.46 | 2.55 | 3.46 | 2.82 |
| Deformed Deviance | 1.06 | | | | | | 0.87 | | | | | |
| Algorithmic Deviance | 0.04 | | | | | | 0.04 | | | | | |
| | 3rd Metatarsal - Lateral Cuneiform | | | | | | Tibia - Talus | | | | | |
| Applied Deformation | 2.00 | 1.00 | 2.00 | -9.00 | 6.00 | -5.00 | 1.00 | -2.00 | 0.00 | -1.00 | 2.00 | -4.00 |
| Deformation in PCA | 2.00 | 1.00 | 2.00 | 11.46 | 6.06 | 11.17 | 1.00 | -2.00 | 0.00 | 1.89 | 4.60 | 4.19 |
| Algorithmic Error | -0.18 | -1.19 | 3.32 | 11.03 | 7.68 | 11.93 | 1.08 | -0.76 | 1.31 | 5.88 | 5.85 | 0.64 |
| Deformed Deviance | 0.45 | | | | | | 0.54 | | | | | |
| Algorithmic Deviance | 0.22 | | | | | | 0.18 | | | | | |

**Table S2** Raw automated alignment results for Case 2. Below each joint, applied random deformation in (x,y,z), applied random deformation in the principle component axes, algorithmic error in principle component axes, normalized deformed deviance, and normalized algorithmic deviance are given. The first three columns for "Applied Deformation" give the applied centroid movement in x, y, and z, respectively, while the final three columns give the applied x, y, and z rotations, respectively. The first three columns in "Deformation in PCA" give the centroid distance between the deformed and ground truth states in x, y, and z, respectively, while the final three columns give the rotations in each principle component axes between the deformed and ground truth states as first major axes, second major axes, and third major axes, respectively. The first three columns in "Algorithmic Error" give the centroid distance between the algorithmic and ground truth states in x, y, and z, respectively, while the final three columns give the rotations in each principle component axes between the algorithmic and ground truth states as first major axes, second major axes, and third major axes, respectively.

| Case #3 | | | | | | | | | | | | |
|---|---|---|---|---|---|---|---|---|---|---|---|---|
| | Medial Cuneiform - Navicular | | | | | | 4th Metatarsal - Cuboid | | | | | |
| Applied Deformation | 0.00 | 2.00 | 2.00 | 8.00 | 3.00 | 8.00 | -2.00 | -2.00 | 1.00 | -6.00 | -2.00 | 0.00 |
| Deformation in PCA | 0.00 | 2.00 | 2.00 | 11.81 | 7.20 | 9.43 | -2.00 | -2.00 | 1.00 | 5.91 | 4.70 | 4.79 |
| Algorithmic Error | -0.35 | -0.35 | 0.84 | 7.55 | 7.32 | 2.32 | -0.63 | -0.46 | 0.26 | 8.46 | 3.83 | 8.74 |
| Deformed Deviance | 0.63 | | | | | | 1.40 | | | | | |
| Algorithmic Deviance | 0.09 | | | | | | 0.01 | | | | | |
| | Intermediate Cuneiform - Navicular | | | | | | 5th Metatarsal - Cuboid | | | | | |
| Applied Deformation | 0.00 | 1.00 | 2.00 | -4.00 | 4.00 | -4.00 | 2.00 | 1.00 | 1.00 | 4.00 | 2.00 | 9.00 |
| Deformation in PCA | 0.00 | 1.00 | 2.00 | 5.88 | 6.67 | 4.37 | 2.00 | 1.00 | 1.00 | 9.51 | 4.73 | 9.56 |
| Algorithmic Error | -0.50 | -0.15 | 0.95 | 0.93 | 0.99 | 1.32 | 4.83 | -2.18 | 0.32 | 17.92 | 8.47 | 18.39 |
| Deformed Deviance | 0.73 | | | | | | 0.22 | | | | | |
| Algorithmic Deviance | 0.04 | | | | | | 0.03 | | | | | |
| | Lateral Cuneiform - Navicular | | | | | | Cuboid - Calcaneus | | | | | |
| Applied Deformation | -2.00 | 2.00 | 0.00 | 8.00 | -9.00 | -6.00 | -2.00 | 2.00 | -2.00 | 10.00 | 10.00 | 9.00 |
| Deformation in PCA | -2.00 | 2.00 | 0.00 | 12.40 | 9.51 | 11.48 | -2.00 | 2.00 | -2.00 | 2.26 | 17.06 | 17.20 |
| Algorithmic Error | -1.29 | 1.71 | -0.87 | 1.83 | 2.68 | 2.88 | 0.39 | 0.14 | 0.04 | 4.90 | 2.25 | 5.10 |
| Deformed Deviance | 0.84 | | | | | | 0.49 | | | | | |
| Algorithmic Deviance | 0.09 | | | | | | 0.10 | | | | | |
| | 1st Metatarsal - Medial Cuneiform | | | | | | Navicular - Talus | | | | | |
| Applied Deformation | -2.00 | 1.00 | -1.00 | 7.00 | 6.00 | 8.00 | 2.00 | -2.00 | 2.00 | 5.00 | 8.00 | -8.00 |
| Deformation in PCA | -2.00 | 1.00 | -1.00 | 10.95 | 10.10 | 9.34 | 2.00 | -2.00 | 2.00 | 8.77 | 8.40 | 12.11 |
| Algorithmic Error | 0.28 | 1.08 | -1.60 | 4.83 | 4.88 | 2.35 | 0.63 | -0.38 | 0.67 | 3.12 | 6.52 | 6.08 |
| Deformed Deviance | 0.27 | | | | | | 0.79 | | | | | |
| Algorithmic Deviance | 0.05 | | | | | | 0.03 | | | | | |
| | 2nd Metatarsal - Intermediate Cuneiform | | | | | | Calcaneus - Talus | | | | | |
| Applied Deformation | 2.00 | -2.00 | 1.00 | -9.00 | -6.00 | -1.00 | 0.00 | 0.00 | 1.00 | 5.00 | 1.00 | -5.00 |
| Deformation in PCA | 2.00 | -2.00 | 1.00 | 9.76 | 6.67 | 9.69 | 0.00 | 0.00 | 1.00 | 6.38 | 7.10 | 3.16 |
| Algorithmic Error | 2.35 | -0.52 | 1.95 | 8.74 | 8.86 | 12.41 | -1.68 | -0.67 | -0.49 | 5.81 | 6.66 | 3.52 |
| Deformed Deviance | 0.87 | | | | | | 0.90 | | | | | |
| Algorithmic Deviance | 0.12 | | | | | | 0.21 | | | | | |
| | 3rd Metatarsal - Lateral Cuneiform | | | | | | Tibia - Talus | | | | | |
| Applied Deformation | 0.00 | 0.00 | -2.00 | -3.00 | -3.00 | 2.00 | -1.00 | -2.00 | 2.00 | 3.00 | 7.00 | 7.00 |
| Deformation in PCA | 0.00 | 0.00 | -2.00 | 4.35 | 1.88 | 4.71 | -1.00 | -2.00 | 2.00 | 7.64 | 9.25 | 8.65 |
| Algorithmic Error | 0.92 | -0.35 | 0.85 | 4.01 | 2.72 | 4.68 | -0.18 | -1.62 | 0.13 | 4.20 | 1.62 | 4.06 |
| Deformed Deviance | 0.76 | | | | | | 0.67 | | | | | |
| Algorithmic Deviance | 0.08 | | | | | | 0.06 | | | | | |

**Table S3** Raw automated alignment results for Case 3. Below each joint, applied random deformation in (x,y,z), applied random deformation in the principle component axes, algorithmic error in principle component axes, normalized deformed deviance, and normalized algorithmic deviance are given. The first three columns for "Applied Deformation" give the applied centroid movement in x, y, and z, respectively, while the final three columns give the applied x, y, and z rotations, respectively. The first three columns in "Deformation in PCA" give the centroid distance between the deformed and ground truth states in x, y, and z, respectively, while the final three columns give the rotations in each principle component axes between the deformed and ground truth states as first major axes, second major axes, and third major axes, respectively. The first three columns in "Algorithmic Error" give the centroid distance between the algorithmic and ground truth states in x, y, and z, respectively, while the final three columns give the rotations in each principle component axes between the algorithmic and ground truth states as first major axes, second major axes, and third major axes, respectively.

| Case #4 | | | | | | | | | | | | |
|---|---|---|---|---|---|---|---|---|---|---|---|---|
| | Medial Cuneiform - Navicular | | | | | | 4th Metatarsal - Cuboid | | | | | |
| Applied Deformation | -2.00 | -2.00 | 2.00 | 7.00 | -3.00 | -8.00 | 2.00 | -2.00 | -1.00 | 10.00 | -2.00 | -3.00 |
| Deformation in PCA | -2.00 | -2.00 | 2.00 | 10.41 | 7.76 | 8.99 | 2.00 | -2.00 | -1.00 | 10.67 | 8.14 | 6.91 |
| Algorithmic Error | -3.00 | -0.14 | 1.14 | 2.07 | 3.84 | 3.86 | 1.24 | 0.88 | -0.75 | 9.84 | 2.98 | 9.41 |
| Deformed Deviance | 0.52 | | | | | | 1.82 | | | | | |
| Algorithmic Deviance | 0.10 | | | | | | 0.27 | | | | | |
| | Intermediate Cuneiform - Navicular | | | | | | 5th Metatarsal - Cuboid | | | | | |
| Applied Deformation | 2.00 | 1.00 | -2.00 | 10.00 | -4.00 | 9.00 | 0.00 | -1.00 | -2.00 | -2.00 | -3.00 | 7.00 |
| Deformation in PCA | 2.00 | 1.00 | -2.00 | 13.51 | 13.21 | 4.85 | 0.00 | -1.00 | -2.00 | 7.69 | 6.21 | 5.25 |
| Algorithmic Error | 2.23 | 0.21 | -2.04 | 9.23 | 9.08 | 12.84 | 0.53 | -0.82 | -0.19 | 3.29 | 7.64 | 8.18 |
| Deformed Deviance | 0.70 | | | | | | 0.96 | | | | | |
| Algorithmic Deviance | 0.08 | | | | | | 0.06 | | | | | |
| | Lateral Cuneiform - Navicular | | | | | | Cuboid - Calcaneus | | | | | |
| Applied Deformation | -1.00 | 2.00 | 1.00 | 8.00 | 0.00 | -9.00 | 2.00 | 1.00 | -2.00 | 9.00 | 3.00 | 9.00 |
| Deformation in PCA | -1.00 | 2.00 | 1.00 | 11.99 | 3.10 | 11.68 | 2.00 | 1.00 | -2.00 | 6.87 | 12.14 | 12.45 |
| Algorithmic Error | -0.42 | 1.99 | -1.45 | 7.69 | 12.41 | 10.47 | 2.49 | 0.33 | -1.26 | 4.44 | 5.86 | 5.77 |
| Deformed Deviance | 0.81 | | | | | | 0.52 | | | | | |
| Algorithmic Deviance | 0.04 | | | | | | 0.07 | | | | | |
| | 1st Metatarsal - Medial Cuneiform | | | | | | Navicular - Talus | | | | | |
| Applied Deformation | -1.00 | -1.00 | 0.00 | 0.00 | 9.00 | 7.00 | 2.00 | 1.00 | 1.00 | -2.00 | -5.00 | 9.00 |
| Deformation in PCA | -1.00 | -1.00 | 0.00 | 1.74 | 11.34 | 11.33 | 2.00 | 1.00 | 1.00 | 8.86 | 6.23 | 10.27 |
| Algorithmic Error | 0.60 | 0.17 | -0.75 | 3.39 | 7.52 | 6.77 | -1.16 | -0.37 | -0.43 | 4.93 | 7.80 | 8.60 |
| Deformed Deviance | 0.67 | | | | | | 0.52 | | | | | |
| Algorithmic Deviance | 0.13 | | | | | | 0.14 | | | | | |
| | 2nd Metatarsal - Intermediate Cuneiform | | | | | | Calcaneus - Talus | | | | | |
| Applied Deformation | -1.00 | 1.00 | -1.00 | -3.00 | 0.00 | -6.00 | -2.00 | -2.00 | -2.00 | 4.00 | 1.00 | 0.00 |
| Deformation in PCA | -1.00 | 1.00 | -1.00 | 5.73 | 4.87 | 5.78 | -2.00 | -2.00 | -2.00 | 3.94 | 2.79 | 3.28 |
| Algorithmic Error | -0.08 | -0.03 | 0.00 | 6.58 | 5.25 | 5.02 | 0.03 | -0.28 | -0.04 | 2.56 | 3.17 | 2.29 |
| Deformed Deviance | 0.80 | | | | | | 0.61 | | | | | |
| Algorithmic Deviance | 0.06 | | | | | | 0.09 | | | | | |
| | 3rd Metatarsal - Lateral Cuneiform | | | | | | Tibia - Talus | | | | | |
| Applied Deformation | -2.00 | 0.00 | -2.00 | -3.00 | 9.00 | 4.00 | -1.00 | 0.00 | 0.00 | -8.00 | -8.00 | -6.00 |
| Deformation in PCA | -2.00 | 0.00 | -2.00 | 3.97 | 10.19 | 9.40 | -1.00 | 0.00 | 0.00 | 12.29 | 9.83 | 8.13 |
| Algorithmic Error | 0.09 | 0.15 | 0.48 | 6.84 | 14.07 | 12.67 | 1.62 | 0.43 | -1.53 | 6.85 | 7.65 | 3.65 |
| Deformed Deviance | 1.09 | | | | | | 0.40 | | | | | |
| Algorithmic Deviance | 0.09 | | | | | | 0.08 | | | | | |

**Table S4** Raw automated alignment results for Case 4. Below each joint, applied random deformation in (x,y,z), applied random deformation in the principle component axes, algorithmic error in principle component axes, normalized deformed deviance, and normalized algorithmic deviance are given. The first three columns for "Applied Deformation" give the applied centroid movement in x, y, and z, respectively, while the final three columns give the applied x, y, and z rotations, respectively. The first three columns in "Deformation in PCA" give the centroid distance between the deformed and ground truth states in x, y, and z, respectively, while the final three columns give the rotations in each principle component axes between the deformed and ground truth states as first major axes, second major axes, and third major axes, respectively. The first three columns in "Algorithmic Error" give the centroid distance between the algorithmic and ground truth states in x, y, and z, respectively, while the final three columns give the rotations in each principle component axes between the algorithmic and ground truth states as first major axes, second major axes, and third major axes, respectively.

| Case #5 | | | | | | | | | | | | |
|---|---|---|---|---|---|---|---|---|---|---|---|---|
| | Medial Cuneiform - Navicular | | | | | | 4th Metatarsal - Cuboid | | | | | |
| Applied Deformation | 0.00 | -1.00 | 1.00 | 2.00 | -9.00 | -3.00 | -1.00 | 1.00 | 0.00 | 2.00 | -6.00 | 9.00 |
| Deformation in PCA | 0.00 | -1.00 | 1.00 | 8.62 | 7.92 | 7.26 | -1.00 | 1.00 | 0.00 | 10.54 | 7.38 | 8.50 |
| Algorithmic Error | -0.90 | -1.37 | -1.97 | 9.70 | 9.07 | 5.75 | -0.47 | 2.76 | 2.46 | 11.86 | 21.76 | 21.24 |
| Deformed Deviance | 0.72 | | | | | | 0.79 | | | | | |
| Algorithmic Deviance | 0.42 | | | | | | 0.16 | | | | | |
| | Intermediate Cuneiform - Navicular | | | | | | 5th Metatarsal - Cuboid | | | | | |
| Applied Deformation | 1.00 | -1.00 | -1.00 | -5.00 | -2.00 | 8.00 | 0.00 | 2.00 | 0.00 | 8.00 | -1.00 | -7.00 |
| Deformation in PCA | 1.00 | -1.00 | -1.00 | 9.03 | 8.99 | 5.12 | 0.00 | 2.00 | 0.00 | 7.52 | 10.18 | 8.33 |
| Algorithmic Error | 0.30 | -1.41 | -0.64 | 3.18 | 4.11 | 5.19 | 1.07 | 0.60 | 1.51 | 6.26 | 2.97 | 6.84 |
| Deformed Deviance | 1.33 | | | | | | 0.44 | | | | | |
| Algorithmic Deviance | 0.03 | | | | | | 0.10 | | | | | |
| | Lateral Cuneiform - Navicular | | | | | | Cuboid - Calcaneus | | | | | |
| Applied Deformation | 2.00 | 1.00 | -2.00 | -6.00 | -2.00 | 9.00 | 0.00 | 2.00 | 1.00 | 1.00 | 4.00 | 8.00 |
| Deformation in PCA | 2.00 | 1.00 | -2.00 | 6.11 | 9.84 | 10.55 | 0.00 | 2.00 | 1.00 | 9.02 | 6.93 | 5.80 |
| Algorithmic Error | 0.42 | -1.36 | -0.55 | 5.18 | 15.23 | 15.71 | -0.97 | -0.11 | -0.18 | 11.34 | 17.19 | 13.42 |
| Deformed Deviance | 1.96 | | | | | | 0.57 | | | | | |
| Algorithmic Deviance | 0.08 | | | | | | 0.53 | | | | | |
| | 1st Metatarsal - Medial Cuneiform | | | | | | Navicular - Talus | | | | | |
| Applied Deformation | 1.00 | -2.00 | -1.00 | 1.00 | -5.00 | 0.00 | -1.00 | 1.00 | 0.00 | -5.00 | 1.00 | -9.00 |
| Deformation in PCA | 1.00 | -2.00 | -1.00 | 4.95 | 4.71 | 2.30 | -1.00 | 1.00 | 0.00 | 10.29 | 4.50 | 9.44 |
| Algorithmic Error | -0.21 | -2.78 | -0.77 | 10.35 | 13.15 | 16.76 | 0.19 | 2.18 | 0.80 | 8.78 | 1.29 | 8.76 |
| Deformed Deviance | 0.85 | | | | | | 0.90 | | | | | |
| Algorithmic Deviance | 0.19 | | | | | | 0.12 | | | | | |
| | 2nd Metatarsal - Intermediate Cuneiform | | | | | | Calcaneus - Talus | | | | | |
| Applied Deformation | -2.00 | 0.00 | 1.00 | 9.00 | 3.00 | 2.00 | 0.00 | 0.00 | -2.00 | 6.00 | -7.00 | 0.00 |
| Deformation in PCA | -2.00 | 0.00 | 1.00 | 6.10 | 8.17 | 9.25 | 0.00 | 0.00 | -2.00 | 8.82 | 8.89 | 3.61 |
| Algorithmic Error | -0.15 | -0.66 | -0.49 | 18.55 | 16.43 | 16.79 | 1.26 | -0.57 | -0.23 | 1.95 | 2.14 | 1.62 |
| Deformed Deviance | 0.39 | | | | | | 0.63 | | | | | |
| Algorithmic Deviance | 0.11 | | | | | | 0.03 | | | | | |
| | 3rd Metatarsal - Lateral Cuneiform | | | | | | Tibia - Talus | | | | | |
| Applied Deformation | 1.00 | 1.00 | -1.00 | 3.00 | -1.00 | 10.00 | 0.00 | 0.00 | 0.00 | 5.00 | 5.00 | -6.00 |
| Deformation in PCA | 1.00 | 1.00 | -1.00 | 10.37 | 8.00 | 6.86 | 0.00 | 0.00 | 0.00 | 7.49 | 7.92 | 6.90 |
| Algorithmic Error | -0.94 | 0.58 | -0.82 | 5.87 | 18.30 | 18.00 | 0.14 | -0.02 | 0.09 | 0.92 | 0.68 | 1.14 |
| Deformed Deviance | 1.02 | | | | | | 0.59 | | | | | |
| Algorithmic Deviance | 0.08 | | | | | | 0.02 | | | | | |

**Table S5** Raw automated alignment results for Case 5. Below each joint, applied random deformation in (x,y,z), applied random deformation in the principle component axes, algorithmic error in principle component axes, normalized deformed deviance, and normalized algorithmic deviance are given. The first three columns for "Applied Deformation" give the applied centroid movement in x, y, and z, respectively, while the final three columns give the applied x, y, and z rotations, respectively. The first three columns in "Deformation in PCA" give the centroid distance between the deformed and ground truth states in x, y, and z, respectively, while the final three columns give the rotations in each principle component axes between the deformed and ground truth states as first major axes, second major axes, and third major axes, respectively. The first three columns in "Algorithmic Error" give the centroid distance between the algorithmic and ground truth states in x, y, and z, respectively, while the final three columns give the rotations in each principle component axes between the algorithmic and ground truth states as first major axes, second major axes, and third major axes, respectively.

| | | | | | | | | | | | | |
|---|---|---|---|---|---|---|---|---|---|---|---|---|
| **Case #6** | | | | | | | | | | | | |
| | Medial Cuneiform - Navicular | | | | | | 4th Metatarsal - Cuboid | | | | | |
| Applied Deformation | 0.00 | 1.00 | 1.00 | -5.00 | -4.00 | 0.00 | -2.00 | 0.00 | -1.00 | -4.00 | 5.00 | 10.00 |
| Deformation in PCA | 0.00 | 1.00 | 1.00 | 3.79 | 5.86 | 5.77 | -2.00 | 0.00 | -1.00 | 10.71 | 6.03 | 11.11 |
| Algorithmic Error | -0.83 | -0.10 | 0.27 | 2.54 | 2.95 | 2.57 | -0.19 | -0.30 | 2.17 | 12.26 | 14.97 | 8.79 |
| Deformed Deviance | 0.98 | | | | | | 0.52 | | | | | |
| Algorithmic Deviance | 0.04 | | | | | | 0.15 | | | | | |
| | Intermediate Cuneiform - Navicular | | | | | | 5th Metatarsal - Cuboid | | | | | |
| Applied Deformation | -1.00 | -1.00 | 2.00 | 5.00 | 7.00 | 0.00 | 0.00 | -2.00 | 0.00 | 2.00 | 7.00 | 9.00 |
| Deformation in PCA | -1.00 | -1.00 | 2.00 | 1.15 | 8.53 | 8.59 | 0.00 | -2.00 | 0.00 | 8.81 | 10.65 | 9.00 |
| Algorithmic Error | -0.03 | -0.18 | 0.40 | 2.75 | 10.12 | 10.28 | 2.18 | -3.09 | -0.83 | 2.65 | 8.73 | 8.44 |
| Deformed Deviance | 1.10 | | | | | | 0.68 | | | | | |
| Algorithmic Deviance | 0.06 | | | | | | 0.10 | | | | | |
| | Lateral Cuneiform - Navicular | | | | | | Cuboid - Calcaneus | | | | | |
| Applied Deformation | -2.00 | 0.00 | 0.00 | 9.00 | 0.00 | 4.00 | -1.00 | -1.00 | 2.00 | 0.00 | 4.00 | -7.00 |
| Deformation in PCA | -2.00 | 0.00 | 0.00 | 8.61 | 8.50 | 6.88 | -1.00 | -1.00 | 2.00 | 7.93 | 2.47 | 7.81 |
| Algorithmic Error | -0.02 | 0.15 | -0.23 | 3.96 | 3.72 | 4.58 | -0.52 | 0.15 | 0.76 | 1.80 | 1.04 | 2.03 |
| Deformed Deviance | 1.07 | | | | | | 0.71 | | | | | |
| Algorithmic Deviance | 0.03 | | | | | | 0.08 | | | | | |
| | 1st Metatarsal - Medial Cuneiform | | | | | | Navicular - Talus | | | | | |
| Applied Deformation | 0.00 | -2.00 | 2.00 | -10.00 | 2.00 | 10.00 | 2.00 | 0.00 | -1.00 | -3.00 | -5.00 | -6.00 |
| Deformation in PCA | 0.00 | -2.00 | 2.00 | 13.19 | 10.31 | 10.94 | 2.00 | 0.00 | -1.00 | 8.25 | 8.26 | 0.70 |
| Algorithmic Error | 3.13 | -0.12 | -1.01 | 17.07 | 9.93 | 18.92 | 1.57 | -0.05 | -0.63 | 6.36 | 1.79 | 6.21 |
| Deformed Deviance | 0.75 | | | | | | 0.39 | | | | | |
| Algorithmic Deviance | 0.17 | | | | | | 0.05 | | | | | |
| | 2nd Metatarsal - Intermediate Cuneiform | | | | | | Calcaneus - Talus | | | | | |
| Applied Deformation | -1.00 | -2.00 | 1.00 | 3.00 | 0.00 | 4.00 | -1.00 | 0.00 | 2.00 | 5.00 | 0.00 | 8.00 |
| Deformation in PCA | -1.00 | -2.00 | 1.00 | 4.45 | 4.42 | 3.26 | -1.00 | 0.00 | 2.00 | 9.38 | 1.00 | 9.43 |
| Algorithmic Error | 1.89 | -0.28 | -0.41 | 2.93 | 2.01 | 3.53 | -0.81 | -0.11 | 0.17 | 4.15 | 3.59 | 2.09 |
| Deformed Deviance | 0.62 | | | | | | 0.85 | | | | | |
| Algorithmic Deviance | 0.10 | | | | | | 0.15 | | | | | |
| | 3rd Metatarsal - Lateral Cuneiform | | | | | | Tibia - Talus | | | | | |
| Applied Deformation | -1.00 | 0.00 | 1.00 | 4.00 | -5.00 | 3.00 | -1.00 | -2.00 | 1.00 | 1.00 | 4.00 | 0.00 |
| Deformation in PCA | -1.00 | 0.00 | 1.00 | 6.63 | 6.98 | 2.25 | -1.00 | -2.00 | 1.00 | 4.12 | 4.12 | 0.27 |
| Algorithmic Error | 1.86 | -0.50 | -0.43 | 1.79 | 1.99 | 1.48 | 0.24 | -0.94 | -0.17 | 4.23 | 1.75 | 3.89 |
| Deformed Deviance | 0.41 | | | | | | 0.53 | | | | | |
| Algorithmic Deviance | 0.07 | | | | | | 0.04 | | | | | |

**Table S6** Raw automated alignment results for Case 6. Below each joint, applied random deformation in (x,y,z), applied random deformation in the principle component axes, algorithmic error in principle component axes, normalized deformed deviance, and normalized algorithmic deviance are given. The first three columns for "Applied Deformation" give the applied centroid movement in x, y, and z, respectively, while the final three columns give the applied x, y, and z rotations, respectively. The first three columns in "Deformation in PCA" give the centroid distance between the deformed and ground truth states in x, y, and z, respectively, while the final three columns give the rotations in each principle component axes between the deformed and ground truth states as first major axes, second major axes, and third major axes, respectively. The first three columns in "Algorithmic Error" give the centroid distance between the algorithmic and ground truth states in x, y, and z, respectively, while the final three columns give the rotations in each principle component axes between the algorithmic and ground truth states as first major axes, second major axes, and third major axes, respectively.

| Case #7 | | | | | | | | | | | | |
|---|---|---|---|---|---|---|---|---|---|---|---|---|
| | Medial Cuneiform - Navicular | | | | | | 4th Metatarsal - Cuboid | | | | | |
| Applied Deformation | 1.00 | -2.00 | -1.00 | 4.00 | -2.00 | 1.00 | -2.00 | -2.00 | 2.00 | -9.00 | 3.00 | -3.00 |
| Deformation in PCA | 1.00 | -2.00 | -1.00 | 2.86 | 4.38 | 3.79 | -2.00 | -2.00 | 2.00 | 9.50 | 9.61 | 4.25 |
| Algorithmic Error | 2.32 | 0.58 | 0.16 | 3.02 | 5.06 | 5.27 | -0.76 | 0.74 | 1.41 | 21.53 | 21.70 | 7.20 |
| Deformed Deviance | 1.11 | | | | | | 1.42 | | | | | |
| Algorithmic Deviance | 0.07 | | | | | | 0.66 | | | | | |
| | Intermediate Cuneiform - Navicular | | | | | | 5th Metatarsal - Cuboid | | | | | |
| Applied Deformation | -2.00 | 1.00 | 2.00 | 1.00 | 2.00 | 3.00 | -2.00 | 1.00 | -2.00 | 8.00 | 1.00 | -4.00 |
| Deformation in PCA | -2.00 | 1.00 | 2.00 | 3.60 | 1.33 | 3.67 | -2.00 | 1.00 | -2.00 | 8.92 | 2.96 | 8.51 |
| Algorithmic Error | -0.32 | -0.41 | 1.34 | 2.13 | 1.97 | 0.82 | -4.66 | -3.11 | -3.42 | 10.39 | 6.67 | 8.02 |
| Deformed Deviance | 1.63 | | | | | | 0.88 | | | | | |
| Algorithmic Deviance | 0.03 | | | | | | 0.08 | | | | | |
| | Lateral Cuneiform - Navicular | | | | | | Cuboid - Calcaneus | | | | | |
| Applied Deformation | -1.00 | -2.00 | 1.00 | 1.00 | 7.00 | 0.00 | 2.00 | 2.00 | 0.00 | -10.00 | 10.00 | 8.00 |
| Deformation in PCA | -1.00 | -2.00 | 1.00 | 6.89 | 5.05 | 5.20 | 2.00 | 2.00 | 0.00 | 2.28 | 15.64 | 15.79 |
| Algorithmic Error | -1.22 | -1.07 | -0.39 | 4.74 | 2.09 | 4.36 | -0.43 | 0.49 | -0.50 | 5.29 | 12.74 | 12.17 |
| Deformed Deviance | 0.58 | | | | | | 0.63 | | | | | |
| Algorithmic Deviance | 0.10 | | | | | | 0.27 | | | | | |
| | 1st Metatarsal - Medial Cuneiform | | | | | | Navicular - Talus | | | | | |
| Applied Deformation | 1.00 | 0.00 | -1.00 | 4.00 | 8.00 | -2.00 | -1.00 | -2.00 | -1.00 | 2.00 | 0.00 | 6.00 |
| Deformation in PCA | 1.00 | 0.00 | -1.00 | 5.06 | 8.53 | 8.21 | -1.00 | -2.00 | -1.00 | 3.90 | 6.07 | 5.29 |
| Algorithmic Error | 0.12 | 0.18 | -0.94 | 1.27 | 3.28 | 3.47 | -1.38 | -0.27 | 0.02 | 5.21 | 2.94 | 5.98 |
| Deformed Deviance | 0.52 | | | | | | 1.17 | | | | | |
| Algorithmic Deviance | 0.06 | | | | | | 0.11 | | | | | |
| | 2nd Metatarsal - Intermediate Cuneiform | | | | | | Calcaneus - Talus | | | | | |
| Applied Deformation | 2.00 | 1.00 | -2.00 | -10.00 | 7.00 | 2.00 | 0.00 | 0.00 | -1.00 | -7.00 | -8.00 | 2.00 |
| Deformation in PCA | 2.00 | 1.00 | -2.00 | 10.02 | 9.31 | 10.65 | 0.00 | 0.00 | -1.00 | 7.86 | 9.38 | 9.36 |
| Algorithmic Error | -0.14 | 1.03 | -1.95 | 4.11 | 6.26 | 4.96 | -1.76 | -0.39 | -0.14 | 2.25 | 5.70 | 6.07 |
| Deformed Deviance | 0.52 | | | | | | 0.87 | | | | | |
| Algorithmic Deviance | 0.12 | | | | | | 0.09 | | | | | |
| | 3rd Metatarsal - Lateral Cuneiform | | | | | | Tibia - Talus | | | | | |
| Applied Deformation | 0.00 | 2.00 | 1.00 | 5.00 | 1.00 | -2.00 | -1.00 | -2.00 | -1.00 | 8.00 | 4.00 | 0.00 |
| Deformation in PCA | 0.00 | 2.00 | 1.00 | 5.46 | 4.94 | 2.32 | -1.00 | -2.00 | -1.00 | 8.86 | 4.19 | 7.99 |
| Algorithmic Error | 1.04 | 1.84 | -3.44 | 6.20 | 11.14 | 10.26 | 0.07 | -0.87 | 0.48 | 10.03 | 2.26 | 10.11 |
| Deformed Deviance | 0.87 | | | | | | 0.57 | | | | | |
| Algorithmic Deviance | 0.16 | | | | | | 0.03 | | | | | |

**Table S7** Raw automated alignment results for Case 7. Below each joint, applied random deformation in (x,y,z), applied random deformation in the principle component axes, algorithmic error in principle component axes, normalized deformed deviance, and normalized algorithmic deviance are given. The first three columns for "Applied Deformation" give the applied centroid movement in x, y, and z, respectively, while the final three columns give the applied x, y, and z rotations, respectively. The first three columns in "Deformation in PCA" give the centroid distance between the deformed and ground truth states in x, y, and z, respectively, while the final three columns give the rotations in each principle component axes between the deformed and ground truth states as first major axes, second major axes, and third major axes, respectively. The first three columns in "Algorithmic Error" give the centroid distance between the algorithmic and ground truth states in x, y, and z, respectively, while the final three columns give the rotations in each principle component axes between the algorithmic and ground truth states as first major axes, second major axes, and third major axes, respectively.

**Case #8**

| | | | | | | | | | | | | |
|---|---|---|---|---|---|---|---|---|---|---|---|---|
| | Medial Cuneiform - Navicular | | | | | | 4th Metatarsal - Cuboid | | | | | |
| Applied Deformation | 2.00 | -1.00 | 0.00 | -8.00 | -10.00 | 8.00 | 1.00 | -1.00 | 1.00 | -10.00 | -1.00 | -8.00 |
| Deformation in PCA | 2.00 | -1.00 | 0.00 | 3.73 | 15.45 | 14.99 | 1.00 | -1.00 | 1.00 | 11.44 | 12.23 | 6.79 |
| Algorithmic Error | -2.00 | 0.70 | -0.45 | 1.97 | 3.18 | 3.59 | 1.08 | -0.55 | 0.02 | 10.57 | 9.68 | 10.01 |
| Deformed Deviance | 0.49 | | | | | | 0.75 | | | | | |
| Algorithmic Deviance | 0.14 | | | | | | 0.25 | | | | | |
| | Intermediate Cuneiform - Navicular | | | | | | 5th Metatarsal - Cuboid | | | | | |
| Applied Deformation | 2.00 | 0.00 | 2.00 | 4.00 | 5.00 | 8.00 | 0.00 | 1.00 | -2.00 | -7.00 | -1.00 | 1.00 |
| Deformation in PCA | 2.00 | 0.00 | 2.00 | 9.92 | 3.15 | 10.35 | 0.00 | 1.00 | -2.00 | 6.46 | 6.53 | 4.23 |
| Algorithmic Error | 1.05 | -0.87 | 0.86 | 8.35 | 4.65 | 9.07 | 2.90 | -1.13 | -3.11 | 3.51 | 3.32 | 4.77 |
| Deformed Deviance | 1.59 | | | | | | 0.84 | | | | | |
| Algorithmic Deviance | 0.03 | | | | | | 0.03 | | | | | |
| | Lateral Cuneiform - Navicular | | | | | | Cuboid - Calcaneus | | | | | |
| Applied Deformation | 1.00 | -1.00 | 2.00 | -6.00 | 0.00 | -4.00 | -1.00 | 1.00 | 0.00 | -3.00 | 8.00 | 2.00 |
| Deformation in PCA | 1.00 | -1.00 | 2.00 | 6.62 | 6.23 | 4.61 | -1.00 | 1.00 | 0.00 | 8.34 | 5.26 | 7.41 |
| Algorithmic Error | 0.54 | -1.53 | 0.95 | 7.46 | 8.13 | 5.09 | -0.01 | -0.51 | 1.20 | 3.04 | 9.20 | 9.50 |
| Deformed Deviance | 1.40 | | | | | | 0.59 | | | | | |
| Algorithmic Deviance | 0.06 | | | | | | 0.07 | | | | | |
| | 1st Metatarsal - Medial Cuneiform | | | | | | Navicular - Talus | | | | | |
| Applied Deformation | 0.00 | 2.00 | 2.00 | -6.00 | -2.00 | -7.00 | -1.00 | -1.00 | 0.00 | -3.00 | 5.00 | 10.00 |
| Deformation in PCA | 0.00 | 2.00 | 2.00 | 8.56 | 6.42 | 7.76 | -1.00 | -1.00 | 0.00 | 9.18 | 168.70 | 7.09 |
| Algorithmic Error | -1.27 | -0.30 | 0.54 | 4.55 | 6.88 | 8.02 | -0.33 | 0.28 | -0.54 | 10.38 | 10.95 | 3.73 |
| Deformed Deviance | 0.77 | | | | | | 1.02 | | | | | |
| Algorithmic Deviance | 0.05 | | | | | | 0.06 | | | | | |
| | 2nd Metatarsal - Intermediate Cuneiform | | | | | | Calcaneus - Talus | | | | | |
| Applied Deformation | -2.00 | -2.00 | -2.00 | 4.00 | 1.00 | -4.00 | -1.00 | 0.00 | 2.00 | 8.00 | -7.00 | 10.00 |
| Deformation in PCA | -2.00 | -2.00 | -2.00 | 5.71 | 3.24 | 4.73 | -1.00 | 0.00 | 2.00 | 12.25 | 8.72 | 13.38 |
| Algorithmic Error | -0.73 | -0.29 | -0.17 | 9.64 | 7.49 | 9.26 | -2.53 | 0.10 | 0.01 | 1.97 | 2.98 | 2.87 |
| Deformed Deviance | 1.73 | | | | | | 1.27 | | | | | |
| Algorithmic Deviance | 0.07 | | | | | | 0.24 | | | | | |
| | 3rd Metatarsal - Lateral Cuneiform | | | | | | Tibia - Talus | | | | | |
| Applied Deformation | 1.00 | 0.00 | 0.00 | 2.00 | -7.00 | -9.00 | 0.00 | -2.00 | -2.00 | -8.00 | 4.00 | 8.00 |
| Deformation in PCA | 1.00 | 0.00 | 0.00 | 7.65 | 8.96 | 11.54 | 0.00 | -2.00 | -2.00 | 8.47 | 8.81 | 11.36 |
| Algorithmic Error | 1.48 | 0.45 | -1.25 | 2.44 | 3.92 | 3.53 | -0.80 | -2.16 | 0.21 | 4.72 | 4.68 | 2.74 |
| Deformed Deviance | 0.55 | | | | | | 0.62 | | | | | |
| Algorithmic Deviance | 0.08 | | | | | | 0.04 | | | | | |

**Table S8** Raw automated alignment results for Case 8. Below each joint, applied random deformation in (x,y,z), applied random deformation in the principle component axes, algorithmic error in principle component axes, normalized deformed deviance, and normalized algorithmic deviance are given. The first three columns for "Applied Deformation" give the applied centroid movement in x, y, and z, respectively, while the final three columns give the applied x, y, and z rotations, respectively. The first three columns in "Deformation in PCA" give the centroid distance between the deformed and ground truth states in x, y, and z, respectively, while the final three columns give the rotations in each principle component axes between the deformed and ground truth states as first major axes, second major axes, and third major axes, respectively. The first three columns in "Algorithmic Error" give the centroid distance between the algorithmic and ground truth states in x, y, and z, respectively, while the final three columns give the rotations in each principle component axes between the algorithmic and ground truth states as first major axes, second major axes, and third major axes, respectively.

| Case #9 | | | | | | | | | | | | |
|---|---|---|---|---|---|---|---|---|---|---|---|---|
| | Medial Cuneiform - Navicular | | | | | | 4th Metatarsal - Cuboid | | | | | |
| Applied Deformation | 0.00 | 1.00 | -1.00 | 5.00 | 9.00 | 8.00 | 0.00 | -1.00 | 1.00 | 6.00 | -5.00 | -3.00 |
| Deformation in PCA | 0.00 | 1.00 | -1.00 | 12.65 | 5.48 | 12.72 | 0.00 | -1.00 | 1.00 | 7.09 | 6.31 | 7.27 |
| Algorithmic Error | -0.10 | 0.59 | -0.64 | 6.26 | 2.82 | 6.82 | -1.65 | 0.71 | -3.14 | 8.53 | 9.55 | 6.27 |
| Deformed Deviance | 0.37 | | | | | | 1.30 | | | | | |
| Algorithmic Deviance | 0.03 | | | | | | 0.11 | | | | | |
| | Intermediate Cuneiform - Navicular | | | | | | 5th Metatarsal - Cuboid | | | | | |
| Applied Deformation | 2.00 | 2.00 | -2.00 | 9.00 | 5.00 | -4.00 | 0.00 | -1.00 | -2.00 | 2.00 | -5.00 | -4.00 |
| Deformation in PCA | 2.00 | 2.00 | -2.00 | 7.31 | 10.76 | 8.25 | 0.00 | -1.00 | -2.00 | 4.81 | 6.71 | 4.82 |
| Algorithmic Error | 0.95 | 0.44 | -1.79 | 10.01 | 7.19 | 10.27 | 2.33 | -0.60 | -1.35 | 5.03 | 2.30 | 5.44 |
| Deformed Deviance | 1.39 | | | | | | 2.00 | | | | | |
| Algorithmic Deviance | 0.04 | | | | | | 0.02 | | | | | |
| | Lateral Cuneiform - Navicular | | | | | | Cuboid - Calcaneus | | | | | |
| Applied Deformation | 0.00 | 1.00 | 1.00 | -1.00 | -9.00 | 4.00 | 0.00 | -1.00 | -2.00 | -1.00 | 4.00 | 4.00 |
| Deformation in PCA | 0.00 | 1.00 | 1.00 | 4.95 | 9.93 | 8.60 | 0.00 | -1.00 | -2.00 | 4.21 | 4.18 | 5.50 |
| Algorithmic Error | 0.34 | -0.33 | -0.22 | 3.24 | 1.34 | 3.11 | 0.74 | 0.83 | -0.93 | 4.12 | 5.63 | 5.67 |
| Deformed Deviance | 1.67 | | | | | | 0.65 | | | | | |
| Algorithmic Deviance | 0.02 | | | | | | 0.10 | | | | | |
| | 1st Metatarsal - Medial Cuneiform | | | | | | Navicular - Talus | | | | | |
| Applied Deformation | 2.00 | 1.00 | 1.00 | 5.00 | 1.00 | -10.00 | -2.00 | 0.00 | 2.00 | 1.00 | 0.00 | -4.00 |
| Deformation in PCA | 2.00 | 1.00 | 1.00 | 10.89 | 5.71 | 9.94 | -2.00 | 0.00 | 2.00 | 3.21 | 3.76 | 3.09 |
| Algorithmic Error | -0.23 | 0.09 | -0.08 | 2.21 | 1.12 | 2.31 | 0.04 | -0.23 | 0.65 | 2.44 | 3.36 | 2.30 |
| Deformed Deviance | 1.00 | | | | | | 1.02 | | | | | |
| Algorithmic Deviance | 0.03 | | | | | | 0.02 | | | | | |
| | 2nd Metatarsal - Intermediate Cuneiform | | | | | | Calcaneus - Talus | | | | | |
| Applied Deformation | 0.00 | -1.00 | 2.00 | 3.00 | 5.00 | -6.00 | 0.00 | -1.00 | -2.00 | -7.00 | -2.00 | -5.00 |
| Deformation in PCA | 0.00 | -1.00 | 2.00 | 8.05 | 1.92 | 8.27 | 0.00 | -1.00 | -2.00 | 8.60 | 3.77 | 8.08 |
| Algorithmic Error | 1.20 | 1.12 | -1.13 | 2.75 | 4.98 | 4.59 | 1.54 | -0.43 | -0.09 | 4.63 | 6.27 | 7.62 |
| Deformed Deviance | 0.77 | | | | | | 1.25 | | | | | |
| Algorithmic Deviance | 0.06 | | | | | | 0.10 | | | | | |
| | 3rd Metatarsal - Lateral Cuneiform | | | | | | Tibia - Talus | | | | | |
| Applied Deformation | -1.00 | 0.00 | 0.00 | 9.00 | 6.00 | 3.00 | 1.00 | -2.00 | 1.00 | -3.00 | 10.00 | 2.00 |
| Deformation in PCA | -1.00 | 0.00 | 0.00 | 9.06 | 9.55 | 9.16 | 1.00 | -2.00 | 1.00 | 10.49 | 10.22 | 3.06 |
| Algorithmic Error | 0.73 | 0.89 | -1.69 | 2.95 | 4.88 | 5.45 | 0.23 | -1.72 | -0.06 | 3.97 | 0.91 | 4.07 |
| Deformed Deviance | 0.71 | | | | | | 0.68 | | | | | |
| Algorithmic Deviance | 0.05 | | | | | | 0.03 | | | | | |

**Table S9** Raw automated alignment results for Case 9. Below each joint, applied random deformation in (x,y,z), applied random deformation in the principle component axes, algorithmic error in principle component axes, normalized deformed deviance, and normalized algorithmic deviance are given. The first three columns for "Applied Deformation" give the applied centroid movement in x, y, and z, respectively, while the final three columns give the applied x, y, and z rotations, respectively. The first three columns in "Deformation in PCA" give the centroid distance between the deformed and ground truth states in x, y, and z, respectively, while the final three columns give the rotations in each principle component axes between the deformed and ground truth states as first major axes, second major axes, and third major axes, respectively. The first three columns in "Algorithmic Error" give the centroid distance between the algorithmic and ground truth states in x, y, and z, respectively, while the final three columns give the rotations in each principle component axes between the algorithmic and ground truth states as first major axes, second major axes, and third major axes, respectively.
